# A Novel CRISPR-Engineered, Stem Cell-Derived Cellular Vaccine

**DOI:** 10.1101/2021.12.28.474336

**Authors:** Krishnendu Chakraborty, Abishek Chandrashekar, Adam Sidaway, Elizabeth Latta, Jingyou Yu, Katherine McMahan, Victoria Giffin, Cordelia Manickam, Kyle Kroll, Matthew Mosher, R. Keith Reeves, Rihab Gam, Elisa Arthofer, Modassir Choudhry, Dan H Barouch, Tom Henley

**Author notes:** Co-corresponding Authors: Tom Henley,; Dan Barouch.

## Abstract

COVID-19 has forced rapid clinical translation of novel vaccine technologies, principally mRNA vaccines, that have resulted in meaningful efficacy and adequate safety in response to the global pandemic. Notwithstanding this success, there remains an opportunity for innovation in vaccine technology to address current limitations and meet the challenges of inevitable future pandemics. We describe a universal vaccine cell (UVC) rationally designed to mimic the natural physiologic immunity induced post viral infection of host cells. Induced pluripotent stem cells were CRISPR engineered to delete MHC-I expression and simultaneously overexpress a NK Ligand adjuvant to increase rapid cellular apoptosis which was hypothesized to enhance viral antigen presentation in the resulting immune microenvironment leading to a protective immune response. Cells were further engineered to express the parental variant WA1/2020 SARS-CoV-2 spike protein as a representative viral antigen prior to irradiation and cryopreservation. The cellular vaccine was then used to immunize non-human primates in a standard 2-dose, IM injected prime + boost vaccination with 1e8 cells per 1 ml dose resulting in robust neutralizing antibody responses (1e3 nAb titers) with decreasing levels at 6 months duration. Similar titers generated in this established NHP model have translated into protective human neutralizing antibody levels in SARS-Cov-2 vaccinated individuals. Animals vaccinated with WA1/2020 spike antigens were subsequently challenged with 1.0 × 10^5^ TCID_50_ infectious Delta (B.1.617.2) SARS-CoV-2 in a heterologous challenge which resulted in an approximately 3-log order decrease in viral RNA load in the lungs. These heterologous viral challenge results reflect the ongoing real-world experience of original variant WA1/2020 spike antigen vaccinated populations exposed to rapidly emerging variants like Delta and now Omicron. This cellular vaccine is designed to be a rapidly scalable cell line with a modular poly-antigenic payload to allow for practical, large-scale clinical manufacturing and use in an evolving viral variant environment. Human clinical translation of the UVC is being actively explored for this and potential future pandemics.

## INTRODUCTION

The COVID-19 pandemic has demonstrated the urgent need for new innovations in vaccinology to enable the rapid development of novel vaccines against emerging viral variants that engender robust and long-lasting immune protection. The unprecedented success of both mRNA and adenoviral vaccines have established the capability of a rapid global vaccination program^1, 2, 3^. However, the waning antibody responses seen with these emergency-use authorized vaccine technologies, and the need for vaccine boosters, has highlighted the requirement for further improvements in vaccine approaches to drive higher, longer-lasting protective immunity^4, 5, 6, 7, 8, 9^. The newly emerging viral variants of SARS-CoV-2, and the evident reduced efficacy of the existing vaccines to protect against transmissible and symptomatic infection of these variants, also highlights the need for vaccines that can ideally deliver multiple variant antigens (polyvalency) and be rapidly manufactured at scale as soon as new viral variants are discovered^10, 11, 12, 13^.

Theoretically, an ideal vaccine technology would have four core attributes, namely: hyper-immunity, self-adjuvancy, polyvalency and scalability. The first hyper-immunity is self-evident and speaks to the requirement of generating a robust humoral neutralizing antibody, and ideally a subsequent T cell amnestic response, such that protection remains durable. Self-adjuvancy, or conversely the absence of the need for exogenous excipients to elicit a hyper-immune response may prove to be a meaningful innovation in that the immune side-effects of current vaccines may be mediated by the non-target antigen specific adjuncts^14^. Thirdly, polyvalency or the ability to protect against multiple immunodominant epitopes, is a core feature of overlapping and orthogonal mechanisms of protection and is a core principle or antibiotic and antiviral therapy in infection to suppress underlying pathogenic genetic drift and mutation and acquired resistance^15, 16^. Lastly, scalability or the ability to deliver preventative doses of vaccines in an economic, large scale and clinically relevant fashion in both the developed and developing worlds, is a *sine qua non* of any human vaccine.

Current mRNA, protein, and viral vector-based vaccines have certain limitations, such as their requirement for excipient adjuvants to activate the recipient immune system, or to deliver the viral antigenic payload^17, 18^. These include the artificial lipid nanoparticles delivering the mRNA, or MF59, AS03, Alum, ISCOMATRIX, and Matrix-M chemical emulsions for example, or the adenoviral protein antigens themselves that stimulate innate immune cell activation^18, 19, 20, 21, 22, 23, 24^. Adjuvants are required to increase the effectiveness of vaccines and their use can cause side-effects including local reactions (redness, swelling, and pain at the injection site) and systemic reactions (fever, chills, and body aches). Furthermore, there have been modest but real morbidity and rare mortality associated with Thrombosis with Thrombocytopenia Syndrome (TTS) and Myocarditis, secondary to the current COVID-19 vaccines^25, 26, 27^.

The size constraint of the adenoviral vector genome, and the limited length of stable mRNA that can be produced and packaged into nanoparticles, restricts the number and size of nucleic acid-encoded antigens and epitopes that can be delivered in these vaccines^28^. Thus, these vaccines are constrained in their ability to provide multiple immunodominant proteins to address emerging pandemic variants, or to easily combine multiple pathogens into one vaccine.

To address some of the current limitations of vaccine technologies, we have developed a novel vaccine platform based on a CRISPR genetically engineered human stem cell, termed the Universal Vaccine Cell (UVC). The principal feature of this vaccine platform is to attempt to reproduce physiologic immunity that is engendered naturally through lytic viral infection and the resulting apoptosis of primary human cells. The platform is designed to deliver an almost unlimited antigenic payload within the context of a physiological apoptotic environment, to both release abundant antigen and simultaneously stimulate the host immune response. Here, we use the SARS-CoV-2 virus as a rigorous and timely test platform, to demonstrate that this self-adjuvanting, polyvalent UVC, can generate a robust and antigen-specific humoral immune response in vaccinated macaques. This hyper-immunogenic vaccine resulted in reduced viral loads in animals challenged with heterologous SARS-CoV-2 variant, which recapitulates the current experience of a population vaccinated against the initial parental WA1/2020 variant, yet now exposed to novel variants like Delta and Omicron^29, 30, 31^.

## RESULTS

### Genetic engineering of iPS cells to create a cellular vaccine to deliver the SARS-CoV-2 spike antigen

To create a cellular vaccine platform to deliver abundant viral antigens and simultaneously engage host innate immune cells to present these antigens to lymphocytes, we attempted to create a cell with a hyper-immunogenic phenotype. We selected human iPS cells as the UVC cell line due to their stable genetics, non-transformed phenotype, ease of genetic engineering and capacity for rapid scalable propagation^32, 33^. IPS cells also retained the unique ability for programmable differentiation into any cell lineage, thus retaining the future opportunity to explore differentiation of the UVC into different cell types that may have enhanced immunogenic properties^34^.

We first genetically engineered iPS cells to create an immunogenic phenotype by stable integration of the SARS-CoV-2 full length spike antigen into the AAVS1 safe-harbor locus using CRISPR/Cas9 gene editing (Fig. 1a). We selected the original and well-characterized WA1/2020 variant of SARS-CoV-2 and spike antigen sequence with mutation of the furin cleavage site and proline-stabilizing mutations that is identical to that in the current emergency-use authorized vaccines being deployed globally to vaccinate against COVID-19^35, 36, 37^ (Supplementary Fig. S1). By including the spike transmembrane domain sequence in the gene encoding this antigen, we were able to detect high levels of the viral spike on the cell surface of the engineered iPS cells (Fig. 1b). Spike protein was also readily observed in engineered cell lysates when measured by western blotting (Fig. 1c). The yield of antigen released upon lysis was quantified using a spike-specific ELISA assay and we observed an abundant and dose-dependent release of protein from the cells, which would equate to approximately ~20 micrograms of spike antigen protein delivered in a 1×10^8^ cell vaccine dose of UVC (Fig. 1d).

**Fig. 1:**
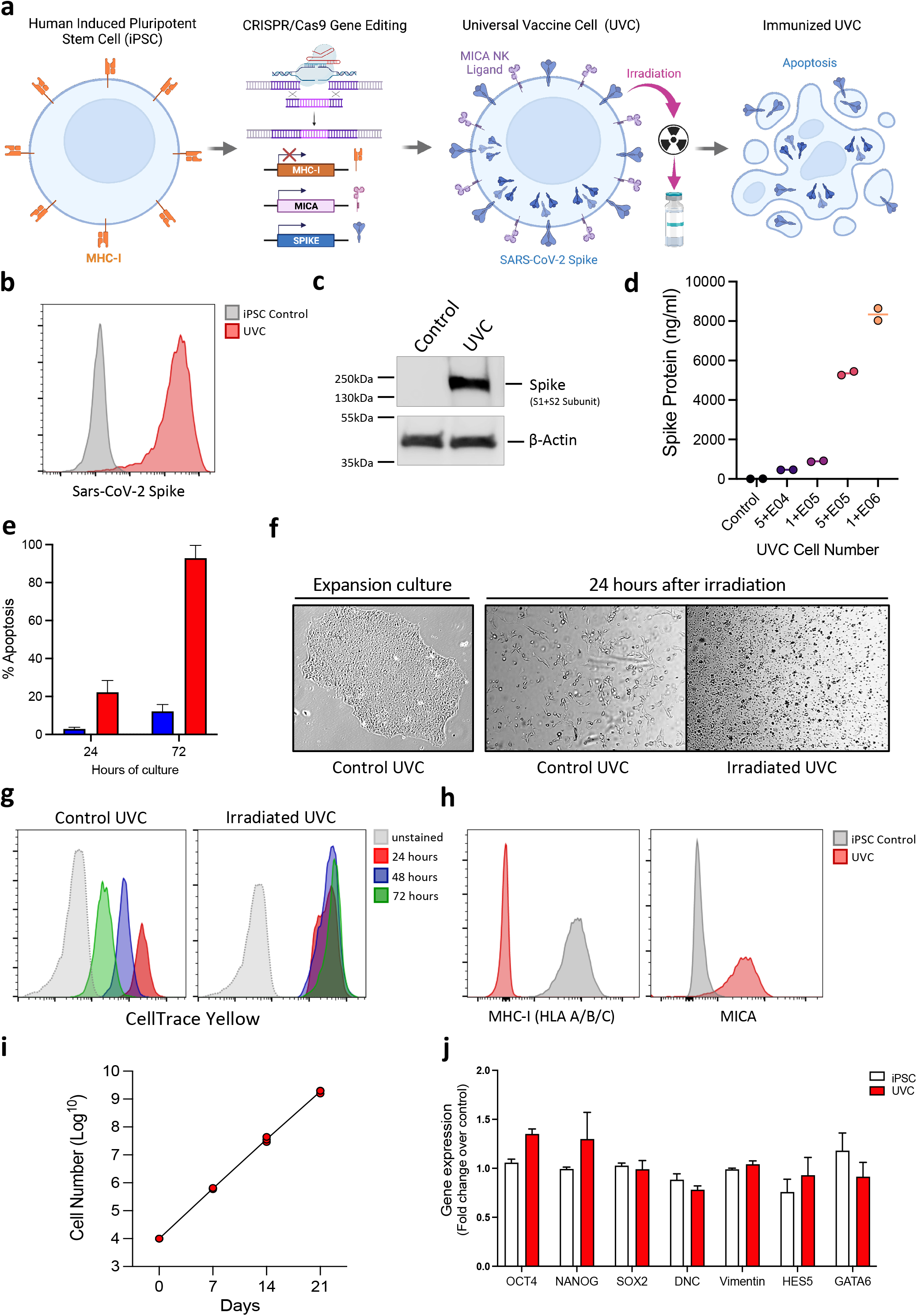
CRISPR genetic engineering of an iPS cell line to create a hyper-immunogenic, self-adjuvanting cellular vaccine. **(a)**, Universal vaccine cell CRISPR genetic engineering strategy to create an apoptotic cellular vehicle for antigen delivery. **(b)**, Representative flow-cytometric analysis showing expression of SARS-CoV-2 WA1/2020 spike protein on the cell surface, and **(c)**, by western blot showing spike protein within UVC whole-cell lysates. **(d)**, ELISA quantification of spike protein released upon UVC lysis. **(e)**, Proportion of apoptotic cells at 24- and 72-hours post-irradiation as measured by 7-AAD staining and flow cytometry. **(f)** Morphology, observed by light microscopy, of engineered UVC during expansion culture, and when reseeded into culture 24 hours after irradiation, showing apoptosis and cell death. **(g)** Absence of detectable proliferation of irradiated UVC as determined by CellTrace Yellow proliferation dye staining and measuring the dilution of the dye by flow cytometry over 72-hours. **(h)** Representative flow-cytometric analysis showing deletion of MHC class-I and overexpression of MICA on the UVC surface by CRISPR engineering. **(i)** Cell counts showing exponential expansion of live engineered UVC over 21-days in culture. **(j)** Relative expression of pluripotency and self-renewal genes by UVC and the control iPS cells from they were derived, as measured by quantitative-PCR, showing maintenance of an iPS cell gene expression prolife after genetic engineering and expansion. Error bars represent mean +/- SEM.

To ensure robust delivery of this immunodominant antigen to the recipient immune system, we incorporate an apoptosis-inducing lethal irradiation step during vaccine manufacture by exposing the UVC cells to a 10 Gy dose of gamma radiation prior to cryopreservation and vaccination. Thus, when subjects are immunized with the UVC, we reasoned that the cells would undergo apoptosis and release the SARS-CoV-2 spike antigen into the immune microenvironment via production of apoptotic bodies (Fig. 1a). In theory, these apoptotic bodies will be phagocytosed by innate immune cells and antigen-presenting cells and presented to T and B lymphocytes to generate a spike antigen-specific immune response.

In addition to creating a mechanism for delivery of immunogenic antigens via apoptotic bodies, the irradiation of the UVC can be considered a safety feature as it renders the cells unable to proliferation or persist *in vivo* upon vaccination. In support of this, we observed a robust elevation in the proportion of apoptotic cells after 24 and 72 hours of culture of irradiated UVC, both using apoptotic dyes and flow cytometry (Fig. 1e) and by observation of cell morphology under the microscope (Fig. 1f). Furthermore, unlike non-irradiated UVC, irradiation prevented any detectible proliferation of the cells over 72-hours in culture as measured by proliferation dyes using flow cytometry (Fig. 1g).

### Incorporation of NK cell activation signals by genetic engineering to create a self-adjuvanting vaccine cell

In addition to the proposed immunogenicity expected from apoptosis and release of immunogenic antigens upon vaccination, we attempted to increase the immunogenic potential further and incorporate a self-adjuvanting phenotype to the UVC. As a form of physiological cell death, apoptosis is generally non-inflammatory^38^. Therefore, to promote effective local inflammation and engage the innate immune system that can mobilize effector cells, we engineered the UVC to mimic a virally infected cell to be recognized and rapidly lysed by host innate immune cells, principally NK cells^39, 40^. Many viruses attempt to evade immune recognition by limiting MHC-I cell surface expression to reduce the presentation of viral antigens to CD8^+^ T cells^41, 42^. This “missing-self’ signal can aid in the activation of NK calls and promote cytolysis, and therefore the iPS cells were engineered to completely remove MHC-I molecules from the cell surface via CRISPR knockout of the β2 microglobulin (B2M) gene, a critical component of MHC class I molecules (Fig. 1h).

*In vivo*, lack of MHC-I on the target cell is not sufficient to trigger full NK cell activation alone and a further hallmark of cells undergoing stress or viral infection, is the expression of NK cell activating natural killer group 2 member D (NKG2D) ligands on their cell surface^43, 44^. Therefore, we further engineered the UVC using CRISPR to integrate a gene expression cassette in a safe-harbor locus to drive constitutive expression of the human MICA gene (MHC class I polypeptide–related sequence A), a potent activator of NK cells. Using flow cytometry, abundant levels of MICA could be detected on the surface of the engineered UVC (Fig. 1h).

### Rapid growth kinetics of engineered UVC

Prior to irradiation and cryopreservation of the UVC ready for immunization, we evaluated the growth kinetics of the cells to confirm the capacity for rapid, scalable proliferation that would be needed for a vaccine technology to address the needs of a pandemic. IPS cells are known to have a relatively short doubling times in the range of 18-20 hours^45, 46^, and we observed similar kinetics with an average exponential growth of >50-fold over a 7-day culture period (Fig. 1i). Thus, from a starting UVC number of 1×10^6^ cells, the vaccine can be theoretically expanded to provide millions of doses in under 8 weeks, and even quicker if adapted to bioreactor manufacturing.

The consistent rapid cell growth of the UVC and the morphological similarity to unmodified iPS cells, suggested the UVC reattained broad characteristic of the iPS cells from which they are derived. We thus assessed the stem cell characteristics of the UVC after genetic engineering and rapid expansion to confirm that the cells have retained their original stemness-gene expression signatures without acquiring any detectible or obvious changes in phenotype beyond those introduced by genetic engineering. The expanded UVC expressing the SARS-CoV-2 spike antigen, human MICA ligand and CRISPR knockout of B2M, showed a similar level of expression of three important pluripotent transcription factors, NANOG, OCT4 and SOX2, suggesting they have retained a stem-cell like transcriptional profile (Fig. 1j). Engineered UVC also showed similar expression to control iPS cells for genes (DNC, Vimentin, HES5 and GATA6) that are known to increase in expression as iPS cells differentiate into mesoderm, endoderm and ectoderm liniage, confirming the UVC have a consistent undifferentiated, iPS cell gene expression profile, morphology, and growth characteristics^47, 48^

Taken together, these data demonstrate that the CRISPR engineered UVC has the capacity to deliver abundant, full-length spike protein antigen in the context of an irradiated, apoptotic cellular vehicle. Via the genetically engineered absence of MHC-I and overexpression of MICA, the UVC has the potential for NK cell activation upon immunization, which may remove the need for excipient adjuvants to promote inflammation and immunogenicity. We first tested this hypothesis *in vitro* to measure the activation of NK cells by the UVC and the NK-mediated UVC cytolysis.

### Human and primate NK cell cytolysis of universal vaccine cells

To further explore the impact of MHC-I loss and overexpression of NK cell ligands on recognition and killing of the UVC by NK cells, we performed a series of *in vitro* NK cell activation and cytolysis assays. When MHC-I was removed via B2M knockout alone, the UVC were robustly killed by human NK cells, which increased in an E:T ratio-dependent manner (Fig. 2a). We compared the level of UVC cytolysis to that observed with the MHC-deficient K562 leukemia cell line, known to be potent targets for NK cell killing, and found a similar level of cytolysis confirming the MHC-I deficient UVC are readily targeted by NK cells. We extended this analysis to macaque NK cells and found that while control iPS cells (expressing MHC-I) show low levels of killing, the MHC-I Knockout UVC were lysed more readily by the NK cells (Fig. 2b).

**Fig. 2:**
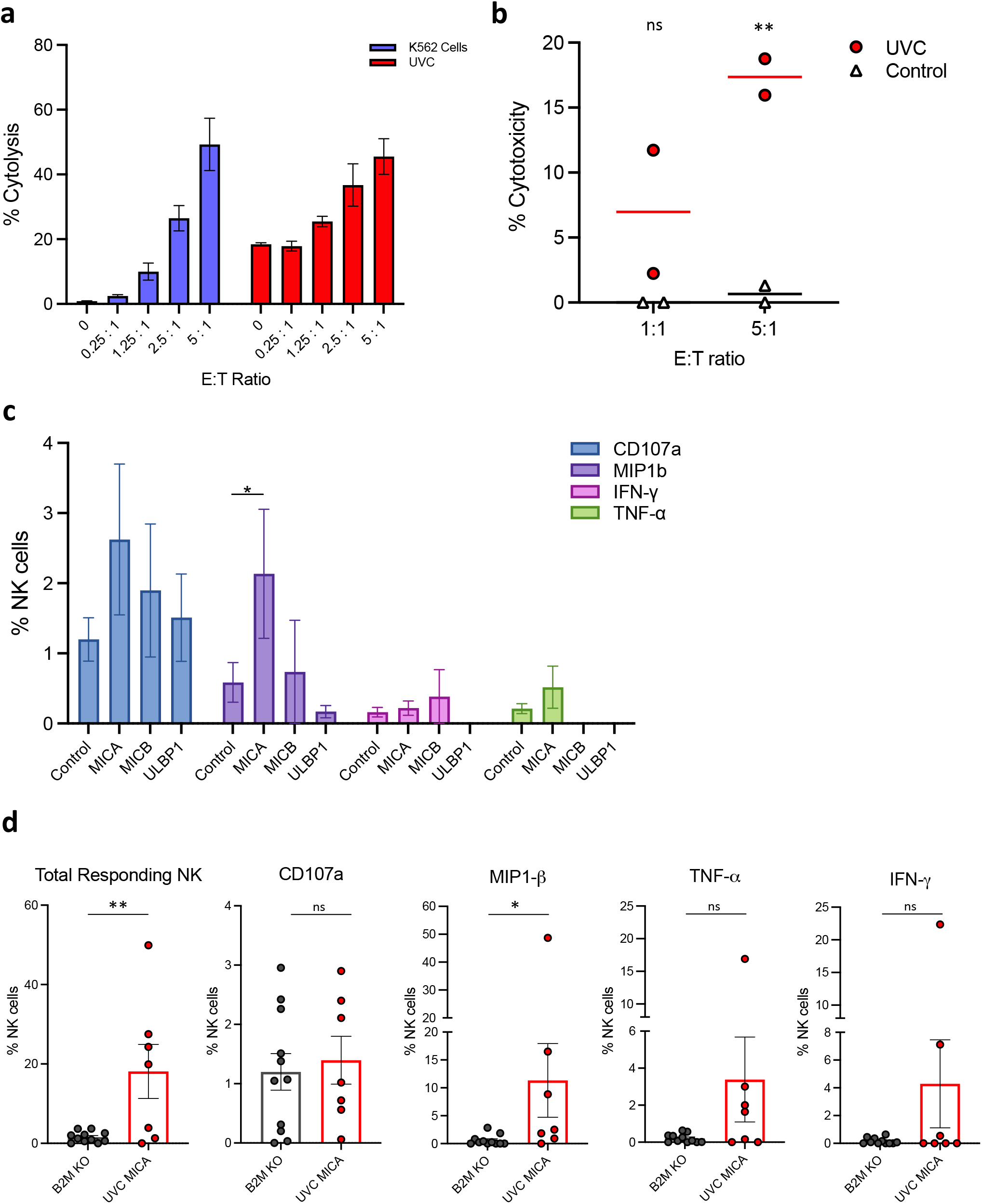
Self-Adjuvancy: Enhanced cytolysis of genetically engineered UVC iPS cells via engineered MHC-I deletion and NK apoptotic ligand expression. **(a)**. CRISPR knockout of B2M and loss of MHC class-I enhances the killing of UVC cells by human primary NK cells, showing equivalent levels of cytolysis seen with the MHC class-I deficient K562 cell line. **(b)**. A similar elevated cytolysis of MHC deficient UVC cells is observed with macaque NK cells. **(c-d)** When overexpressed transiently on the UVC, NKG2D family ligands show no elevation in markers of NK cell activation by macaque NK cells, except MICA which significantly elevates levels of macrophage inflammatory protein-1β (MIP-1β). **(d)** When stably overexpressed on the UVC by CRISPR editing, MICA enhanced the NK cell functional responses as measured by ICS. **P<0.01, Error bars represent mean +/- SEM.

To assess the relative contribution of overexpressing NK activating ligands on UVC cytolysis by macaque NK cells, we performed an analysis of UVC cells transiently overexpressing different NKG2D ligands, including MICA, MICB and UL16 binding protein 1 (ULBP1). While the levels of macrophage inflammatory protein-1β (MIP-1β) was significantly elevated when MICA was overexpressed, proinflammatory and activation markers for NK cells were generally the same regardless of ligand overexpression (Fig. 2c). With stable overexpression of MICA by CRISPR engineering, we confirmed a significant increase in total responding macaque NK cells and a significant elevation in MIP-1β (Fig. 2d).

Collectively these data demonstrate robust cytolysis of the UVC by both human and non-human primate NK cells and show the potent impact of loss of MHC-I on NK cell-mediated lysis. We hypothesized that overexpression of the NK cell activating ligand MICA, in combination with loss of MHC-I, would lead to robust recognition and activation of NK cells *in vivo* upon immunization and enable delivery of the vaccine antigen and engender a potent immune response in the absence of any adjuvant. We thus moved to non-human primate immunization studies to investigate further.

### Immunogenicity of universal vaccine cells in vaccinated macaques

To evaluate the immunogenicity of the UVC and the vaccine’s ability to engender a humoral immune response, we immunized cynomolgus macaques and followed the production of neutralizing and spike-specific antibodies over a 10-week period, and a duration follow up at 6 months. We immunized 9 macaques, aged 6-12 years old, with either 1×10^7^ UVC (*n*=3) or 1×10^8^ UVC (*n*=3) expressing the WA1/2020 SARS-COV-2 spike antigen, and sham controls (*n*=3). Macaques received a prime dose immunization by the intramuscular route without adjuvant at week 0, followed by a boost dose immunization (same cell number as prime dose) at week 6 (Fig. 3a). Neutralizing antibody responses were assessed using a pseudovirus neutralization assay^49, 50, 51, 52^, and we observed neutralizing antibodies in all UVC vaccinated macaques at week 2 that further increased by week 4 (Fig. 3b). The higher dose of 1×10^8^ UVC resulted in the most robust titers of neutralizing antibodies at all timepoints tested. Following UVC boost dose immunization at week 6, neutralizing antibody titers elevated further, reaching close to 1×10^3^ titers with the higher 1×10^8^ cell dose. Six months after the initial UVC immunization, neutralizing antibody showed a durable response, and levels in macaques immunized with the 1×10^8^ UVC dose remained elevated beyond that seen with the initial prime UVC dose. We also observed robust spike-specific and receptor-binding-domain (RBD)-specific antibody titers, as measured by enzyme-linked immunosorbent assay (ELISA) in vaccinated macaques (Fig. 3c and d). These antibody responses and durability at 6 months in the 1×10^8^ dose, were like those seen with the neutralizing antibody titers, and thus collectively demonstrating that the UVC vaccine can engender a robust humoral response against the SARS-CoV-2 spike antigen with demonstrable durability. At 6 months after immunization, detectible levels of neutralizing antibodies against Beta and Delta were also observed, albeit lower than seen with the immunizing antigen variant WA1/2020 spike, suggesting humoral immunity is also generated against SARS-CoV-2 variants (Fig. 3e). This prompted us to assess the protective immunity of the humoral immune response generated by UVC immunization in the context of a SARS-CoV-2 heterologous infection challenge.

**Fig. 3:**
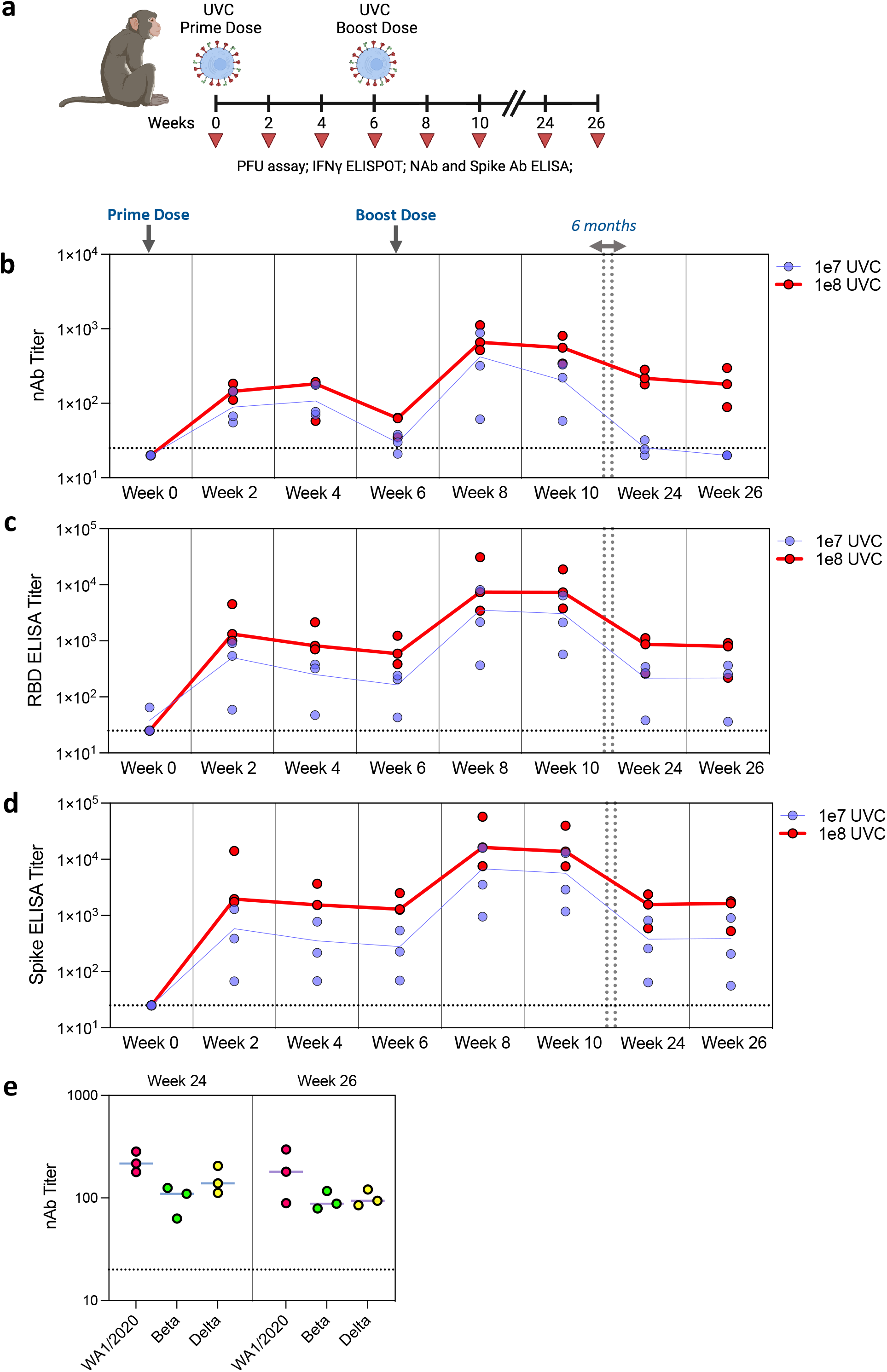
Humoral immune responses in UVC vaccinated macaques. **(a)** Macaques received a high WA1/2020 spike expressing UVC prime dose (1×10^8^) or low UVC prime dose (1×10^7^) at week 0, and a boost dose matched to that of the prime dose at week 6. Humoral immune responses were assessed at 2-week intervals up to week-10 and then again at weeks 24 and 26 by **(b)** spike, **(c)** RBD-specific binding antibody ELISA, and **(d)** pseudovirus neutralization assays. **(e)** In addition to the WA1/2020 SARS-CoV-2 variant, detectible neutralizing antibodies against the B.1.351 (Beta) and B.1.617.2 (Delta) variants were observed in immunized macaques at weeks 24 and 26. Red bars reflect median responses. Dotted lines reflect assay limit of quantification. NAb, neutralizing antibody.

### Humoral immune responses in vaccinated macaques after heterologous SARS-CoV-2 challenge

In a second non-human primate study, we immunized rhesus macaques, aged 6-12 years old, with the higher 1×10^8^ dose of UVC (*n*=6) expressing the SARS-CoV-2 WA1/2020 spike antigen, and sham controls (*n*=6), and this time followed the production of neutralizing and spike-specific antibodies over an 8-week period (Fig. 4a). At week 8, the macaques were challenged with 1.0 × 10^5^ 50% tissue culture infectious dose (TCID_50_) of heterologous SARS-CoV-2 B.1.617.2 (Delta) by the intranasal and intratracheal routes^51, 52^. Viral loads in bronchoalveolar lavage (BAL) and nasal swabs were assessed over 10-days by reverse transcription PCR (RT–PCR) specific for subgenomic mRNA (sgRNA), which is thought to measure replicating virus^52, 53^. Sham controls showed a median peak of 5.39 (range 4.60– 5.88) log10[sgRNA (copies per ml)] in BAL samples (Fig. 4b and d). Partial protection was observed in macaques immunized with the UVC as a significantly lower level of virus was detected in BAL samples, with a median peak of 2.78 (range 1.70–4.63) log10[sgRNA (copies per ml)], representing a 2.81 log reduction in virus in UVC vaccinated animals. A significant reduction in virus (0.96 log reduction) was also observed in nasal swabs from UVC immunized macaques when compared to sham controls, albeit lower in magnitude than seen when comparing BAL samples (Fig. 4c and e).

**Fig. 4:**
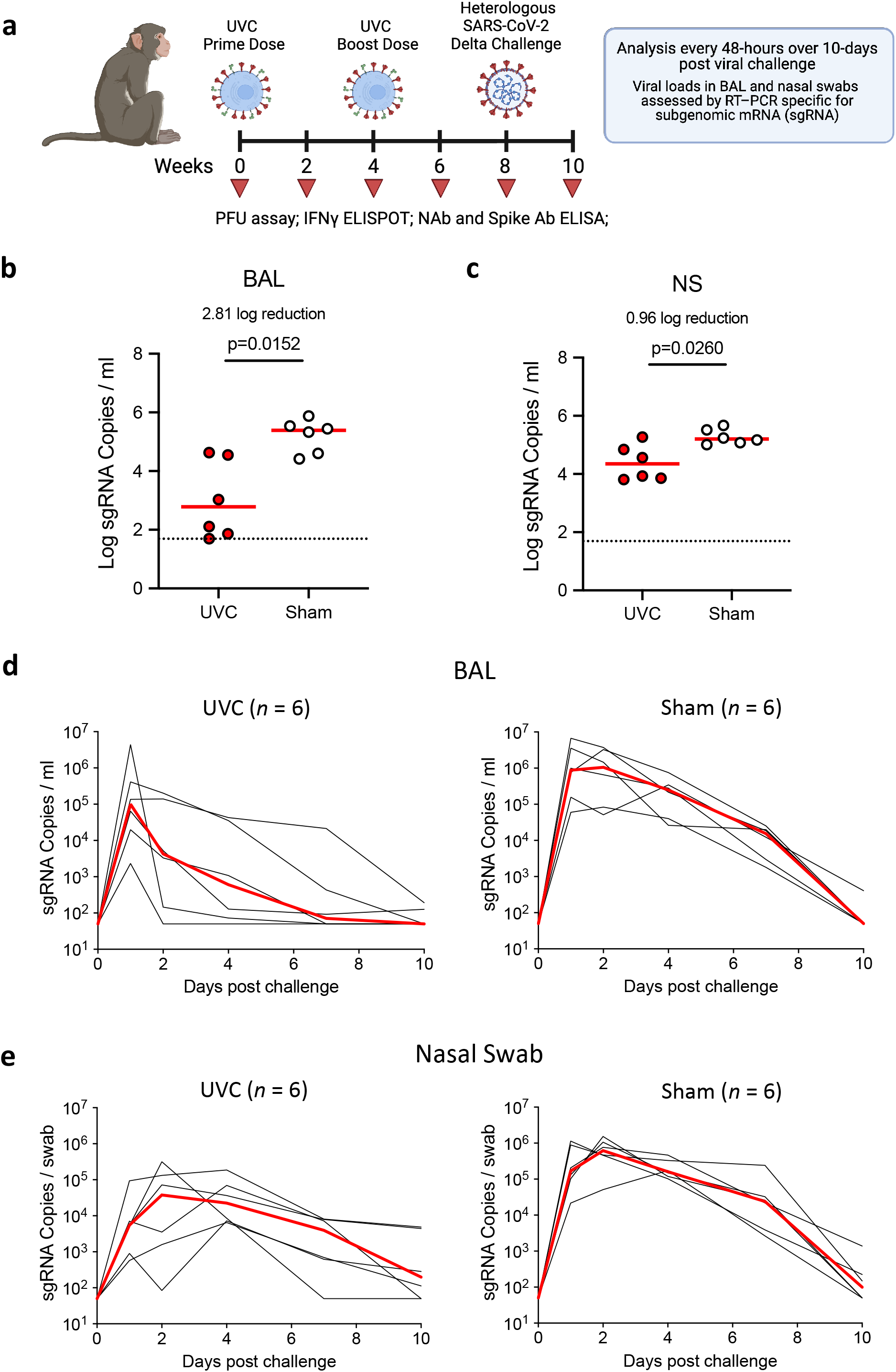
Viral loads in UVC vaccinated macaques after heterologous SARS-CoV-2 challenge. **(a)** Rhesus macaques were immunized with 1×10^8^ WA1/2020 spike expressing UVC at week 0 and received a boost dose of 1×10^8^ matched UVC at week 4. Macaques were then challenged at week 6 by the intranasal and intratracheal routes with 1.0 × 10^5^ TCID_50_ of SARS-CoV-2 B.1.617.2 (Delta). Log10[sgRNA (copies per ml)] (limit of quantification 50 copies per ml) were assessed, and peak viral loads are shown in **(b)** bronchoalveolar lavage (BAL) samples, and **(c)** Nasal swabs (NS), in sham controls and vaccinated macaques after challenge. Viral loads were assessed every 2 days **(d-e)**. Dotted lines reflect assay limit of quantification. NAb, neutralizing antibody.

While neutralizing antibody titers specific for the WA1/2020 variant spike was high in both macaque immunization studies (Fig. 3 and supplementary Fig. S2), the titers specific for other SARS-CoV-2 variants (Beta and Delta) was lower, which is to be expected given the divergence in antigen protein sequence. Thus, the partial protection seen in animals immunized with WA1/2020 spike UVC and challenged with B.1.617.2 (Delta) SARS-CoV-2, is also expected given the heterologous nature of the challenge. We would predict a more robust and complete reduction in virus from animals immunized with UVC and challenged with SARS-CoV-2 in which the antigen and variant are matched.

Collectively these data demonstrate that a prime and boost dose of 1×10^8^ WA1/2020 Spike expressing UVC promote a robust antigen-specific antibody response with levels of neutralizing antibodies and durability similar to the current approved COVID-19 vaccines^49, 54, 55^, and this can lead to partial protective immunity in a heterologous WA1/2020 versus Delta SARS-CoV-2 virus challenge.

## DISCUSSION

The urgent global need for improved vaccine technologies to meet future pandemics is driving a renaissance of innovation in vaccinology. COVID-19 has demonstrated the rapid pace at which viral mutations can accumulate and new variants emerge that can escape the protective efficacy of existing vaccines designed against earlier viral antigen sequences^10, 11, 29, 30, 31, 56, 57^. To address the need for novel vaccine technologies, we have developed the first cellular vaccine, to generate hyper-immunity via self-adjuvancy through apoptosis and NK cell-mediated cytolysis within the immune microenvironment.

Our data demonstrate that the UVC vaccine platform can induce robust neutralizing antibody responses in vaccinated macaques when delivering the SARS-CoV-2 WA1/2020 (original variant^58^) full-length membrane-bound Spike protein, with mutation of the furin cleavage site and two proline-stabilizing mutations^35, 36, 37^. The antibody titers and resulting immunogenicity were on par with the neutralizing antibody titers demonstrating immune protection with the emergency use authorized and now commercially available mRNA vaccines (1e3 nAb titer)^49,54,55^. We also show that the UVC delivering the WA1/2020 Spike antigen enhances neutralizing antibodies and (RBD)-specific binding antibodies specific for the B.1.617.2 (Delta) and B.1.351 (Beta) variants of SARS-CoV-2. This robust and specific humoral response can partially protect rhesus macaques vaccinated with a prime and boost dose of WA1/2020 Spike UVC in a *heterologous* variant challenge with infectious B.1.617.2 (Delta) SARS-CoV-2 and engender a more rapid clearance of viral RNA in the BAL. A limitation of the WA1/2020 UVC in a Delta heterologous challenge was the absence of any meaningful protection in nasal swabs. This discordance between vaccine variant, versus virus variant, is the principal challenge facing current vaccinated populations and might suggest the reason for ongoing infectious spread but more limited morbidity and mortality against these emerging variants^29, 31^.

As regards duration of protection, the theoretical hyper-immunity postulated by creating a self-adjuvanting, hyper-immune UVC established robust initial nAb titers. When these animals were rechallenged 6-month later, the nAb response stayed robust at the higher, and now established for clinical use, 1e8 UVC dose. Moreover, the persistent nAb response at 6-months remained robust for SARS-CoV-2 WA1/2020, Beta and Delta variants.

As regards intrinsic safety, the UVC undergoes lethal irradiation during manufacture and rapid apoptosis in the immune microenvironment upon vaccination. This is the principal mechanism of efficacy of the UVC, and fortunately its most redeeming safety feature, by virtue of the impossibility of *in vivo* persistence and teratogenicity of the cellular antigen carrier. The irradiation-induced apoptosis is further enhanced by CRISPR genetic engineering to remove MHC-I expression and introduce cell surface expression of the NKG2D ligand MICA, making the UVC potent targets for host NK cells. Recruited NK cells will likely recognize the UVC as virally infected cell through MHC-I absence and MICA activation of NKG2D signaling to mediate a direct killing effect and release of protein antigen^59^. The apoptosis and NK-mediated cytolysis enables the UVC to be a self-adjuvanting vaccine vector, without the need for additional chemicals adjuvants or additional foreign antigens. Thus, the UVC may mimic the physiological engagement of the immune system typical of virally infective cells within the tissues of an individual suffering with the disease. The use of a living cellular antigen vehicle, as opposed to a lipid nanoparticle or other such inanimate construct, can potentially recapitulate natural immunity without the need for exogenous adjuvants, which may portend greater safety against autoimmune complications. Theoretically, and unproven and unprovable at this time absent human clinical data, the natural physiologic nature may lead to an improved safety profile for the UVC.

The CRISPR genetic engineering to render the UVC highly immunogenic and self-adjuvanting, also presents a unique opportunity to address antigen polyvalency. Unlike mRNA or DNA vaccines, or recombinant, replication-incompetent viral vector vaccines, that have a size limit of encoded antigen or the number of independent antigens they can deliver, the UVC can be engineered to express and deliver a much higher number of full-length protein antigens. Thus, there is the ability to create polyvalency against multiple epitopes in a rapid modular gene cassette fashion through CRISPR engineering of the iPS cell genome.

A perceived limitation of the UVC technology may be the seemingly complex and costly nature of developing and manufacturing a live human cell as a vector for vaccination at scale. However, the UVC is a cell line, not a complex cell therapy, and can thus be scaled within appropriate parameters for such a biologic agent. Once pathogen antigens have be genetically engineered, the UVC cell line can be expanded rapidly to scale with predictable growth kinetics and QA/QC controls. The modular nature of the UVC and the ability to integrate emerging viral antigens into the cellular genome using CRISPR, can allow scalable manufacture of new polyvalent vaccines to address emerging variants. In fact, the genetic engineering of the UVC cells can be accomplished in a matter of weeks prior to exponential cell culture expansion to create millions of clinical doses. As a test of the rapid and modular manufacturing of the platform, at the time of this publication, the authors have begun engineering a polyvalent UVC against the SARS-CoV-2 Omicron variant (B.1.1.529) (Fig. 5b).

**Fig. 5:**
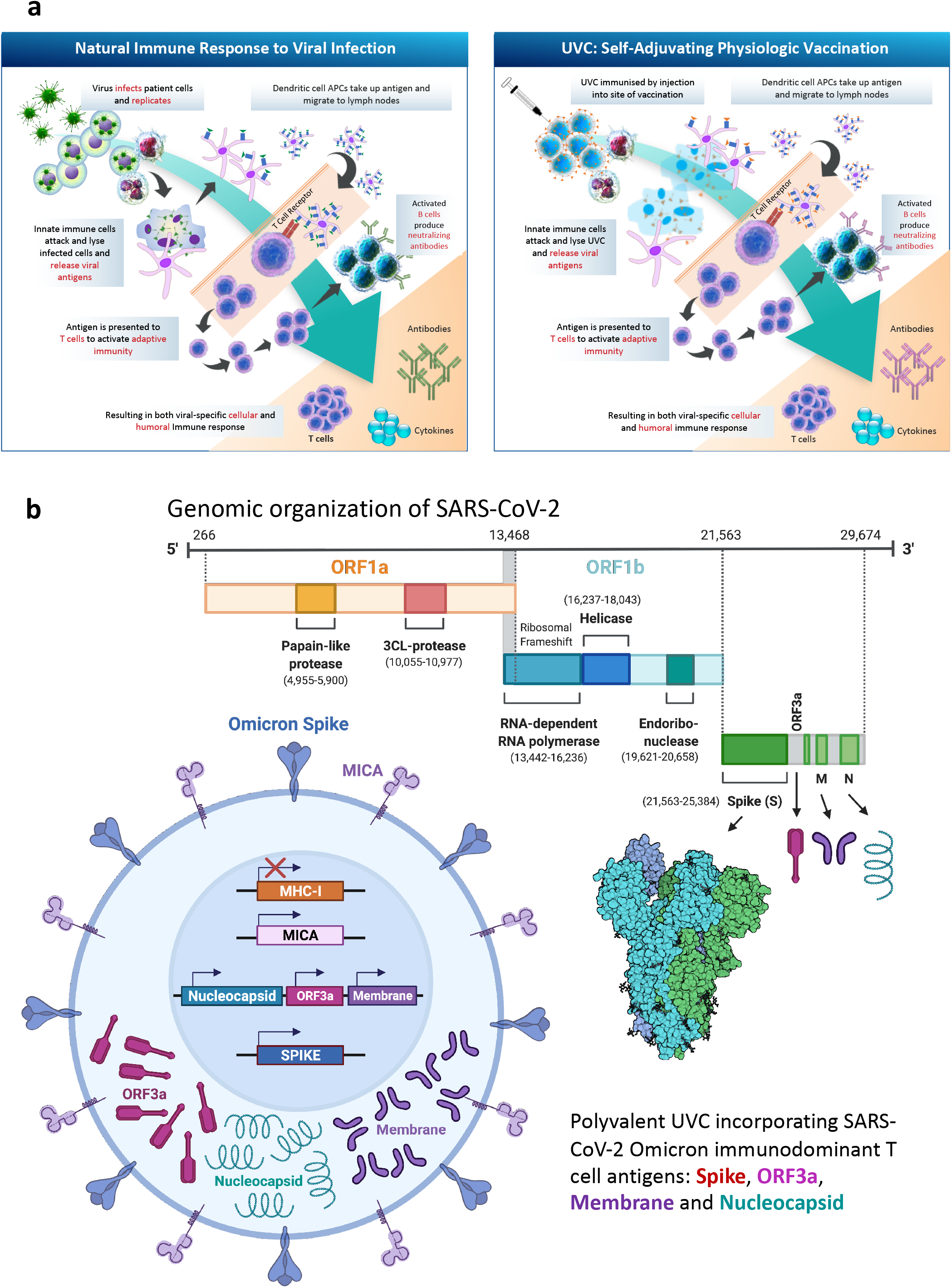
(a) Proposed mechanism of action of UVC to replicate natural physiologic immunity and (b) polyvalent UVC design for emerging variants (Omicron). Schematic depiction of the UVC design to deliver a polyvalent SARS-Cov-2 B.1.1.529 Omicron variant vaccine, incorporating the Spike, Nucleocapsid, ORF3a and Membrane viral proteins

A limitation of our study is that we have yet to observe the generation of a robust T cell response in vaccinated animals. To put this into broader clinical context, the measurable albeit modest CD8-specific T cell responses seen with the adenoviral and mRNA vaccines have not resulted in nAb titers and duration of protection longer than 6-9 months^8, 60^. One potential hypothesis to begin to establish a clinically meaningful amnestic T cell response is to explore novel non-spike antigens, thus leveraging the simultaneous polyvalency of the UVC platform. The UVC allows vaccination with multiple SARS-CoV-2 antigens, including immunodominant T cell epitopes such as those in the nucleocapsid and viral accessory proteins^61, 62, 63^. In the context of COVID-19, additional studies will evaluate the capability of UVCs delivering polyvalent T cell epitopes such as the Nucleocapsid, ORF and Membrane proteins, to generate a CD8 cytolytic and CD4 Helper T cell response in vaccinated macaques to engender duration of protection via T cell amnestic response. Beyond COVID-19, the potential of the theoretically unlimited antigenic payload of a cellular vaccine would allow for “poly-pathogenic polyvalency” and the creation of a single seasonal respiratory vaccine, which could include influenza, RSV, and pan-coronavirus viral antigens^64^.

In summary, these data establish the first cellular vaccine platform and demonstrate that immunization with a WA1/2020 SARS-CoV-2 spike expressing UVC vaccine elicits robust neutralizing antibody titers and provides partial protection against heterologous Delta SARS-CoV-2 challenge in rhesus macaques. Establishing this novel cellular vaccine platform technology within the rigorous and timely setting of COVID-19, the UVC may portend a novel class of gene and cell therapy prophylaxis for potential future viral pandemics.

## METHODS

### iPS cell culture, irradiation, and cryopreservation

Human iPS cells (Thermo Fisher) were cultured on vitronectin-coated T225cm^2^ flasks using complete mTesSR Plus medium (StemCell Technologies) supplemented with 1% penicillin/streptomycin, Rock inhibitor (StemCell Technologies) at 1:1000 dilution. For drug selection, G148 was used at 500ug/ml and puromycin at 5ug/ml (Sigma-Aldrich). Cultures were maintained at 37 °C, 5% CO2 in a humidified incubator. Harvesting of engineered UVC was performed using accutase (StemCell Technologies) and cells were counted using a CellDrop cell counter (DeNovix). Cells were irradiated at a total single dose of 10 Gy, before centrifugation at 300 xg for 10 minutes followed by resuspension in 100 μl of CryoStor-CS10 freezing media (StemCell Technologies). The UVC preparations for use in non-human primate studies were analyzed for endotoxin levels (Wickham Laboratories Ltd) and absence of mycoplasma (Mycoplasma Experience Ltd).

### CRISPR genetic engineering

CRISPR sgRNAs targeting the human B2M gene, PPP1R12C (AAVS1), and the ROSAβgeo26 locus were designed and validated for indel formation at the selected genomic site. Up to 6 sgRNAs per target gene were tested and the most efficient sgRNA was selected containing 2’-O-methyl and 3’ phosphorothioate modifications to the first three 5’ and the last three 3’ nucleotides (Synthego). 2×10^6^ UVC cells were electroporated using a Neon Nucleofector (Lonza) in Buffer P3 (Lonza) with Cas9 protein (IDT) precomplexed with sgRNA, in a total volume of 100 μl using electroporation program CM138. Gene targeting vectors carrying an expression cassette for expression of human MICA or the SARS-CoV-2 WA1/2020 spike gene, targeting the Rosa26 and AAVS1 locus respectively, were co-electroporated at 4 μg. Indels introduced by CRISPR editing were detected by PCR and Sanger sequence using DNA primers designed to amplify a 600-900 base pair region surrounding the sgRNA target site. A minimum of 24 hours after electroporation, genomic DNA was extracted using the DirectPCR Lysis solution (Viagen Biotech) containing Proteinase K and target regions were amplified by PCR using the GoTaq G2 PCR mastermix (Promega). Correct and unique amplification of the target regions was verified by agarose gel electrophoresis before purifying PCR products using the QIAquick PCR Purification Kit (Qiagen). For analysis by TIDE, PCR amplicons were Sanger sequenced (Eurofins or Genewiz) and paired .ab1 files of control versus edited samples were analyzed using Synthego’s ICE tool (https://ice.synthego.com).

### Intracellular spike protein staining

Engineered UVC were harvested and then fixed and permeabilized using BD Cytofix/Cytoperm Fixation/Permeabilization Solution (ThermoFisher). Cells were then stained for intracellular spike protein using an Anti-SARS-CoV-2 Spike Glycoprotein S1 antibody (Abcam, ab275759, 1:50) followed by Goat Anti-Rabbit IgG H&L (Alexa Fluor 488) (Abcam, ab150077, 1:500). Flow analysis was carried out on a Fortessa flow cytometer (BD Bioscience), and data analyzed, and flow cytometry figures generated using FlowJo 10 software (BD Biosciences).

### Flow cytometry analysis of cell surface antigen expression

For flow cytometric analysis of cell surface expression of MHC-I, MICA and SARS-CoV-2 spike protein, cells were harvested from culture plates and washed using PBS with 1% Bovine Serum Albumen (Thermo Scientific) and were then stained with PE anti-human MICA/MICB Antibody (6D4, Biolegend), Alexa Fluor 647 anti-human HLA-A,B,C (W6/32, Biolegend), and anti-SARS-CoV-2 Spike Glycoprotein S1 antibody (Abcam, ab275759, 1:50) followed by Goat Anti-Rabbit IgG H&L (Alexa Fluor 488) (Abcam, ab150077, 1:500). Live/Dead Fixable Dead Cell Stains (Invitrogen) were included in all experiments to exclude dead cells. After staining, cells were resuspended in PBS with 2% Human Heat Inactivated AB Serum (Sigma) and 0.1 M EDTA pH 8.0 (Invitrogen) before analysis on a Fortessa flow cytometer (BD Bioscience) and data analyzed using FlowJo 10 software (BD Biosciences).

### Western blot

The SARS-CoV-2 spike glycoprotein was detected in UVC lysates by western blotting. Briefly, cells were lysed by RIPA buffer (20 mM Tris-HCl pH 7.5, 150 mM NaCl, 1 mM EDTA, 0.1% SDS, 1% NP40, 1x protease inhibitor cocktail). Samples were spun at 4°C for 10 mins at 12,000 xg and the pellet discarded. Protein content was measured using BCA Assay (ThermoFisher) using a PHERAstar plate reader (BMG Labtech) at 560 nm. LDS Sample Buffer was added to 30 ng of protein sample to make a 1x solution, with 0.5 μl of b-mercaptoethanol per well and heated at 70°C for 10 minutes before separation on a polyacrylamide gel (Bio-Rad Mini-PROTEAN TGX Gel 4-15%) and transferred to a PVDF membrane. Membranes were blocked in blocking buffer (5% non-fat powdered milk in TBST), before incubation with primary antibodies in blocking buffer (Rabbit polyclonal anti-SARS-Cov2, Sino Biological 40591-T62, 1:6000 dilution or Mouse b-actin, Abcam 8226, 1 μg/ml), detected with HRP conjugated secondaries in blocking buffer (Goat anti-Rabbit HRP, Sino Biological SSA003, 0.5 μg/ml or Goat anti-Mouse HRP, Abcam ab205719, 1: 4000 dilution) and visualised using the SuperSignal West Femto kit (ThermoFisher) as per kit instructions.

### qPCR measurement of stem cell factors

Total RNA was extracted from UVC cells using the ReliaPrep RNA miniprep (Promega) according to the manufacturer’s instructions (a DNase treatment was included for all samples), and RNA concentration and absorbance ratios were measured using a Nanodrop One Spectrophotometer (ThermoFisher). cDNA was synthesized using a High-Capacity cDNA Reverse Transcription Kit (the Applied Biosystems) in a total volume of 20 μl to produce DNA that was subsequently assessed by spectrophotometric analysis and diluted to 100 ng/μl. Individual master mixes with each of the DNA-primer combinations for detection of human SOX2, NANOG, OCT4, DNC, Vimentin, HES5 and GATA6 genes were made for 3 replicates using the Brilliant III Ultra-Fast SYBR green qPCR master mix (Agilent Technologies) and analyzed on a CFX Opus Real-Time PCR system (BioRad) using the following program: 95 °C for 15 minutes for 1 cycle; 95°C for 15 seconds for 40 cycles; 60°C for 30 seconds.

### SARS-CoV-2 spike protein ELISA

Cell pellets were harvested and lysed in 20 μl Cell Extraction Buffer (Invitrogen) containing protease inhibitors (Sigma) on ice for 30 minutes, with 3 brief vortexing every 10 minutes. Samples were centrifuged at 13,000 rpm for 10 minutes at 4°C to pellet insoluble contents. S1 Spike protein was detected using a Covid-19 S-protein ELISA kit (Abcam) specific to S1RBD. Samples were diluted to a range determined to be within the working range of the ELISA kit used and the assay procedure was follows as per manufacturer’s instructions. The resulting colorimetric signal was detected at 450 nm using a PHERAstar (BMG LABTECH) plate reader. GraphPad Prism was used to plot a standard curve and interpolate the sample values using a 4-parameter logistic fit.

### UVC Proliferation and apoptosis assays

To quantify apoptosis of UVC post-irradiation, cells were stained using a FITC Annexin V Apoptosis Detection Kit with 7-AAD (Biolegend). Proliferation of cells was measured staining of control and Irradiated UVC with either 2 μM Cell Trace Yellow (Abcam) according to kit protocol and analyzing the dilution of the dye at 24-hour periods over 3-days and measuring fluorescence intensity. Flow analysis was carried out on a Fortessa flow cytometer (BD Bioscience), and data analyzed, and flow cytometry figures generated using FlowJo 10 software (BD Biosciences).

### CAM cytotoxicity assay

Both MHC-I expressing and MHC-I deficient (B2M knockout) UVC were used as target cells for NK cell cytotoxicity assay. Trypsinized cells were stained with calcein acetoxymethyl ester (CAM, Invitrogen) at a 10 μM concentration for 1 hour at 37°C and then washed to remove excess dye. NK cells highly enriched from normal cynomolgus macaque (*Macaca fascicularis*) blood samples using a CD3 depletion kit (Miltenyi Biotec), were used as effector cells. NK cell effectors and stained target cells were co-cultured in 96 well round bottom plates at effector: target (E:T) ratios of 1:1 and 5:1. Control wells included – only target cells for spontaneous release of CAM and target cells treated with Triton-X100 for maximum release of CAM. At the end of 4-hour incubation, supernatant was collected for CAM measurement in a fluorescent plate reader at 530 nm. Percent-specific lysis = (test release - spontaneous release)/(maximum release - spontaneous release).

### Nucleofection of NKG2D ligands in iPS cells

UVC were cultured in EGM2 (Lonza) media supplemented with 20 ng/ml VEG-F (Peprotech) until 70-90% confluent, in tissue culture flasks pre-coated with sterile 0.1% gelatin in PBS for 1 hour at 37°C. The cells were removed from culture flasks using trypsin, washed, and transfected with plasmid DNA containing either MICA, MICB or ULBP-1 genes after optimizing nucleofection conditions using primary cell 4D nucleofector kit and 4D nucleofector system (Lonza). After 48 hours of culture, transfected cells were stained with aqua dye for live/dead discrimination and corresponding antibodies-MICA/MICB (Clone 6D4, PE, BioLegend) or ULBP-1 (clone 170818, PE, R & D Systems). Stained cells were fixed with 2% paraformaldehyde and acquired on LSRII flow cytometer. Transfection efficiency was calculated as % live cells expressing transfected protein.

### NK cell intracellular cytokine staining assay

NK cell effectors were enriched from normal cynomolgus macaque (Macaca fascicularis) blood samples using a CD3 depletion Kit (Miltenyi Biotec). Target and effector cells were plated at E:T ratio of 2:1 in a 96 well round bottom plate. Anti-CD107a antibody (clone H4A3, ECD conjugate, BD Biosciences), brefeldin A and monensin (BD Biosciences) were added to all the samples prior to incubation. After 6 hours of incubation at 37°C, the cells were washed and stained with aqua dye used for live and dead cell discrimination for 20 minutes at room temperature. The cells were then washed and stained for surface markers that included CD3 (SP34.2, BV421, BD Biosciences), CD14 (M5E2, BV650, BD Biosciences), CD16 (3G8, BUV496, BD Biosciences), CD20 (L27, BV570, BD Biosciences), CD56 (NCAM1.2, BV605, BD Biosciences), HLA-DR (G46-6, APC-H7, BD Biosciences) and NKG2A (Z199, PE-Cy7, BD Biosciences) to delineate NK effector cells. Following incubation for 20 minutes, cells were washed and permeabilized using fix & perm reagent (Thermofisher Scientific) as per manufacturer’s recommendation. Intracellular cytokine staining was performed for macrophage inflammatory protein 1β (MIP-1β; D21-1351, FITC, BD Biosciences) interferon-γ (IFN-γ; B27, BUV395, BD Biosciences), tumor necrosis factor-α (TNF-α; Mab11, BV650, BD Biosciences) at 4°C for 15 minutes.

Cells were washed, fixed, and acquired on LSRII flow cytometer. Unstimulated NK cells were used for background subtraction of percent positive cells. NK cells stimulated with leukocyte activation cocktail (BD Biosciences) were used as positive control for the assay.

### Animals and study design

Outbred adult male and female rhesus macaques (*M. mulatta*) and cynomolgus macaques (*M. fascicularis*), 6–12 years old, were randomly allocated to groups. All macaques were housed at Bioqual. Macaques were treated with irradiated UVC at doses of either 1×10^7^ or 1×10^8^ cells (*n* = 3-6), and sham controls (*n* = 3-6). Prior to immunization, the cryopreserved doses of irradiated UVC were thawed at 37°C, then 900 μl of 1xPBS was added to each vial of 100 μl UVC in CryoStore freezing media. Macaques received a prime immunization of 1ml of UVC by the intramuscular route without adjuvant at week 0. At weeks 4 or 6, macaques received a boost immunization of either 1×10^7^ or 1×10^8^ UVC. At week 10 all macaques were challenged with 1.0 × 10^5^ TCID_50_ (1.2 × 10^8^ RNA copies, 1.1 × 10^4^ PFU) SARS-CoV-2, which was derived from B.1.617.2 (Delta). Viral particle titers were assessed by RT– PCR. Virus was administered as 1 ml by the intranasal route (0.5 ml in each nare) and 1 ml by the intratracheal route. All immunological and virological assays were performed blinded. All animal studies were conducted in compliance with all relevant local, state, and federal regulations and were approved by the Bioqual Institutional Animal Care and Use Committee (IACUC).

### Subgenomic viral mRNA assay

SARS-CoV-2 *E* gene sgRNA was assessed by RT–PCR using primers and probes as previously described^49, 50, 51, 52^. In brief, to generate a standard curve, the SARS-CoV-2 *E* gene sgRNA was cloned into a pcDNA3.1 expression plasmid; this insert was transcribed using an AmpliCap-Max T7 High Yield Message Maker Kit (Cellscript) to obtain RNA for standards. Before RT–PCR, samples collected from challenged macaques or standards were reverse-transcribed using Superscript III VILO (Invitrogen) according to the manufacturer’s instructions. A Taqman custom gene expression assay (ThermoFisher Scientific) was designed using the sequences targeting the *E* gene sgRNA. Reactions were carried out on a QuantStudio 6 and 7 Flex Real-Time PCR System (Applied Biosystems) according to the manufacturer’s specifications. Standard curves were used to calculate sgRNA in copies per ml or per swab; the quantitative assay sensitivity was 50 copies per ml or per swab.

### Serum antibody ELISA

RBD-specific binding antibodies were assessed by ELISA as previously described^9,10^. In brief, 96-well plates were coated with 1 μg ml-1 SARS-CoV-2 RBD protein (A. Schmidt, MassCPR) in 1× DPBS and incubated at 4 °C overnight. After incubation, plates were washed once with wash buffer (0.05% Tween 20 in 1× DPBS) and blocked with 350 μl casein block per well for 2–3 hour at room temperature. After incubation, block solution was discarded, and plates were blotted dry. Serial dilutions of heat-inactivated serum diluted in casein block were added to wells and plates were incubated for 1 hour at room temperature, before three further washes and a 1-hour incubation with a 1:1,000 dilution of anti-macaque IgG HRP (NIH NHP Reagent Program) at room temperature in the dark. Plates were then washed three times, and 100 μl of SeraCare KPL TMB SureBlue Start solution was added to each well; plate development was halted by the addition of 100 μl SeraCare KPL TMB Stop solution per well. The absorbance at 450 nm was recorded using a VersaMax or Omega microplate reader. ELISA endpoint titers were defined as the highest reciprocal serum dilution that yielded an absorbance >0.2. The log_10_(endpoint titers) are reported.

### Pseudovirus neutralization assay

The SARS-CoV-2 pseudovirus expressing a luciferase reporter gene were generated in a similar approach to that previously described^9,10,16^. In brief, the packaging construct psPAX2 (AIDS Resource and Reagent Program), luciferase reporter plasmid pLenti-CMV Puro-Luc (Addgene), and spike protein expressing pcDNA3.1-SARS-CoV-2 SΔCT were co-transfected into HEK293T cells with calcium phosphate. The supernatants containing the pseudotype viruses were collected 48 hours after transfection; pseudotype viruses were purified by filtration with 0.45-μm filter. To determine the neutralization activity of the antisera from vaccinated macaques, HEK293T-hACE2 cells were seeded in 96-well tissue culture plates at a density of 1.75 × 10^4^ cells per well overnight. Twofold serial dilutions of heat-inactivated serum samples were prepared and mixed with 50 μl of pseudovirus. The mixture was incubated at 37 °C for 1 hour before adding to HEK293T-hACE2 cells. After 48 hours, cells were lysed in Steady-Glo Luciferase Assay (Promega) according to the manufacturer’s instructions. SARS-CoV-2 neutralization titers were defined as the sample dilution at which a 50% reduction in relative light units was observed relative to the average of the virus control wells.

### Statistical Analyses

Statistical differences between two sample groups, where appropriate, were analyzed by a standard Student’s two-tailed, non-paired, t-test and between three or more sample groups using two-way or three-way ANOVA using GraphPad Prism 9. Analysis of virological data was performed using two-sided Mann–Whitney tests. Correlations were assessed by two-sided Spearman rank-correlation tests. P values are included in the figures or referred to in the legends where statistical analyses have been carried out. P values of less than 0.05 were considered significant.

## ETHICS DECLARATION

D.B. has a sponsored research collaboration funded by Intima Bioscience. Praesidium Bioscience has patents filed based on the findings described herein.

## Supplementary

**(S1).**
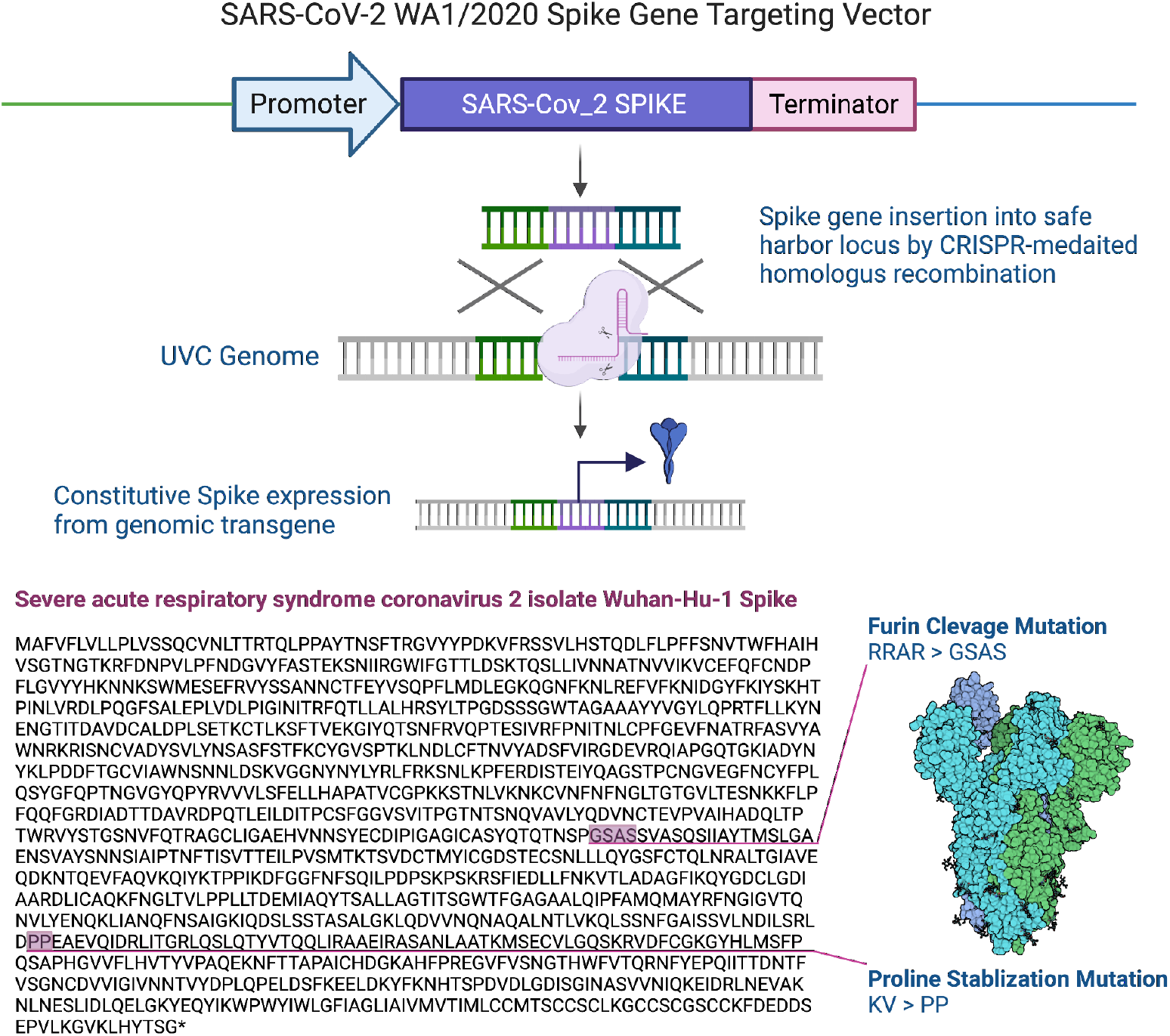
Design of the gene targeting vector to insert the SARS-CoV-2 spike gene into the UVC genome by CRISPR-mediated homology-directed repair. The amino acid sequence of the WA1/2020 Spike protein is shown highlighting the furin cleavage and proline-stabilizing mutations.

**(S2).**
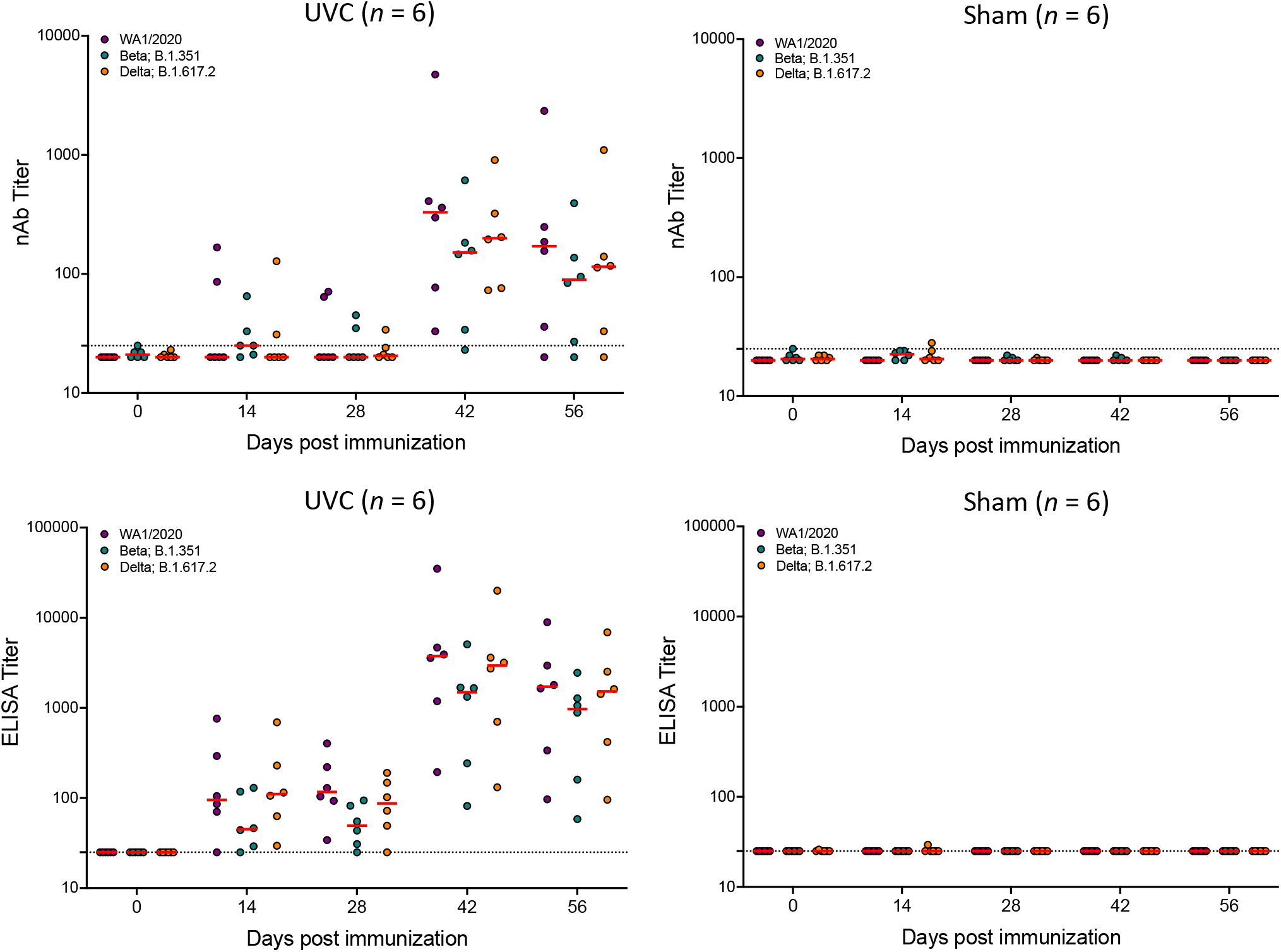
Humoral immune responses in WA1/2020 spike expressing UVC vaccinated rhesus macaques prior to challenge with SARS-CoV-2 B.1.617.2 (Delta). Antibody responses were assessed at weeks 0, 2, 4, 6 and 8 by pseudovirus neutralization assay and spike specific antibody ELISA. Red bars reflect median responses. Dotted lines reflect assay limit of quantification. NAb, neutralizing antibody.

## Notes

### Competing Interest Statement

The authors have declared no competing interest.

